# Higher-Order Interactions in Auxotroph Communities Enhance Their Resilience to Resource Fluctuations

**DOI:** 10.1101/2024.05.22.595348

**Authors:** Tong Wang, Ashish B. George, Sergei Maslov

**Affiliations:** Department of Biological Sciences, Purdue University, West Lafayette, IN 47907, USA; Carl R. Woese Institute for Genomic Biology, University of Illinois at Urbana-Champaign, Urbana, IL 61801, USA; Department of Plant Biology, University of Illinois at Urbana-Champaign, Urbana, IL 61801, USA; Broad Institute of MIT and Harvard, Cambridge, MA 02142, USA; Department of Bioengineering, University of Illinois at Urbana-Champaign, Urbana, IL 61801, USA

## Abstract

Auxotrophs are prevalent in microbial communities, enhancing their diversity and stability—a counterintuitive effect considering their dependence on essential resources from other species. To address the ecological roles of auxotrophs, our study introduced a novel consumer-resource model that captures the complex higher-order interactions within these communities. We also developed an intuitive graphical and algebraic framework, which assesses the feasibility of auxotroph communities and their stability under resource fluctuations and biological invasions. Validated against experimental data from synthetic *E. coli* auxotroph communities, the model accurately predicted outcomes of community assembly. Our findings highlight the critical role of higher-order interactions and resource dependencies in maintaining the diversity and stability of microbial ecosystems dominated by auxotrophs.

## INTRODUCTION

Auxotrophs, organisms that cannot synthesize essential resources and are therefore forced to rely on their production by other species, are extremely common in microbial communities [1–4]. Despite the vulnerability of individual auxotrophic species to essential resource deprivation, research shows that communities with high proportions of auxotrophs exhibit increased diversity and stability [5]. Previous studies have investigated this phenomenon through different approaches, including experimental work on obligate cross-feeding in synthetic microbial communities [6] and a stochastic single-cell model of gene expression and growth dynamics [7]. However, a mechanistic, community-level model that quantitatively captures the ecological benefits of auxotrophy remains lacking and is critically needed.

Traditional models, which assume that growth rates of auxotrophs depend linearly on the abundances of other species that synthesize missing essential resources, do not adequately capture the complex dynamics within auxotroph communities [8, 9]. For example, a generalized Lotka-Volterra (gLV) model, which focuses on such pairwise interactions, completely failed to predict the assembly outcome of a community of 14 *E. coli* synthetic auxotrophic strains [10]. In fact, it predicted the coexistence of all 14 strains, while only 4 strains survived in experiments. MacArthur’s Consumer-Resource Model (CRM), which accounts only for resource consumption and microbial growth, cannot capture the dynamics of resource exchange between auxotrophs [11]. However, when CRM is extended to include resource production, cross-feeding, or multiple essential resources, it has been shown to exhibit higher-order interactions when mapped to the species-species modeling framework [12]. The growth of auxotrophs is described by Liebig’s law of minimum, which in turn depends on the concentrations of multiple essential resources (e.g., 20 amino acids). In this case, the growth rate of one auxotrophic species may depend not just on the abundance of a single species, but on the presence of multiple species that collectively determine its limiting resource. This results in higher-order interactions between species [13, 14] and underscores the need for a more sophisticated modeling approach than the pairwise gLV model.

In this study, we use the CRM framework with resource exchanges [11, 15, 16], which more accurately represents resource concentrations and acquisition mechanisms. We introduce and study a novel consumer-resource model of auxotroph communities that goes beyond simple pairwise interactions and captures essential higher-order interactions and resource dependencies. Each microbial species in our model is capable of transforming the primary resource (e.g., glucose) into multiple essential resources (e.g., amino acids), a capability that auxotrophs lack for some resources. Thus, our model can account for both auxotrophic and prototrophic species.

We also develop an intuitive graphical and algebraic framework that assesses the feasibility of auxotroph communities and their stability under resource fluctuations and biological invasions by other species. Our approach offers insights into complex interactions and resource strategies in auxotroph communities. Its success is highlighted by a successful prediction of 3 out of 4 surviving strains and extinction of the remaining 10 strains in communities of *E. coli* synthetic auxotrophic strains.

## MODEL AND RESULTS

### Consumer Resource Model of Auxotroph/Prototroph Microbial Communities

We aim to model a general situation in which microbial species are capable of transforming (green arrows in Fig. 1a) a single externally supplied carbon source (e.g., glucose) into multiple essential resources (e.g., amino acids). We assume that a fraction of essential resources is stoichiometrically transformed to biomass (red arrows in Fig. 1a), while the rest overflows into the environment (orange arrows in Fig. 1a). We also assume that all species can freely use overflow resources released by other species (black arrows in Fig. 1a). Lastly, we assume that the stoichiometry in which essential resources are used for biomass synthesis is the same across all species.

**FIG. 1.**
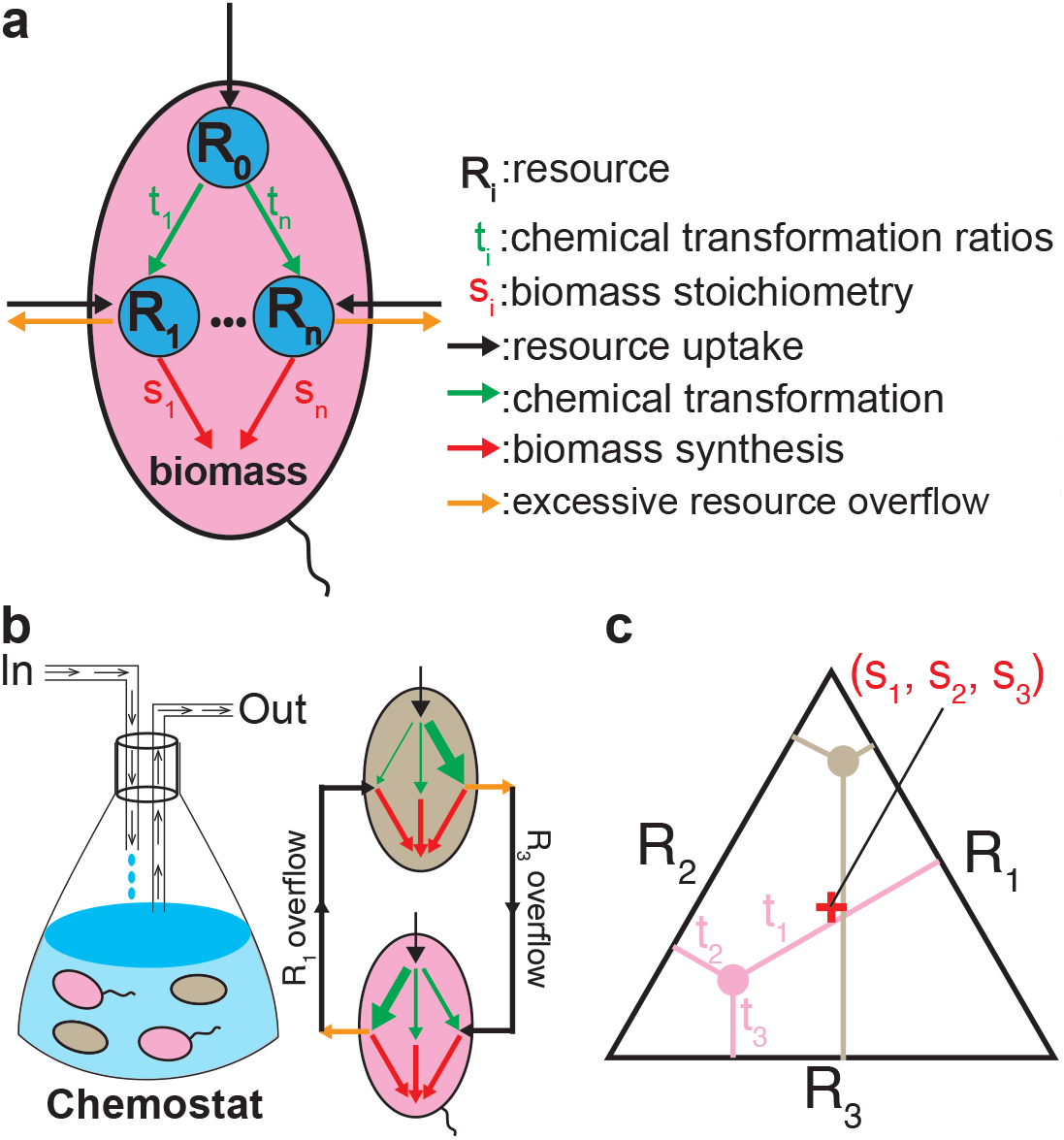
Conversion and exchange of resources in our model of auxotroph communities. **a** Schematic illustration of intracellular processes in one microbial species. Each species is capable of transforming a primary resource *R*_0_ (e.g., glucose) into a subset of other resources (*R*_1_,…,*R*_*n*_) essential for growth (e.g., amino acids). Different species have unique transformation vectors 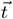 quantifying this conversion but share the same biomass stoichiometry 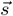. If the transformation flux of an essential resource (green arrows) exceeds its flux directed to biomass (red arrows), the excess is secreted as the overflow flux (orange arrows). Otherwise, a microbe can uptake essential resources directly from the environment (black arrows) to balance its biomass synthesis flux. **b** Depiction of a microbial community in a chemostat (red arrows). The pink microbe uptakes *R*_3_ and produces *R*_1_ due to the excessive transformation of *R*_1_. The green-grey microbe uptakes *R*_1_ and produces *R*_3_. As a result, two microbial species cooperate via the exchange of essential metabolites. **c** Graphical representation of the microbial community (panel b) in the resource simplex when 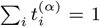. Two species, represented as pink and brown dots inside a simplex, have identical biomass stoichiometry 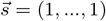 but different transformation fluxes. The distances from a species’ position inside a simplex to its three sides are proportional to its corresponding transformation fluxes (*t*_1_, *t*_2_, and *t*_3_). The red cross at the simplex’s center represents the shared stoichiometry vector.

In mathematical notation, the fluxes transforming one unit of the primary resource (green arrows in Fig. 1a) into multiple essential resources (*R*_*i*_ with *i* = 1 : *n* in Fig. 1a) are represented as a transformation vector 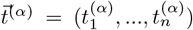 (green arrows in Fig. 1a). Thus the transformation flux at which one species *α* generates essential resource *i* is modeled as 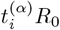, where *R*_0_ is the concentration of the primary resource. All species share the same biomass synthesis stoichiometry, denoted by *s* = (*s*_1_, …, *s*_*n*_) (red arrows in Fig. 1a). This assumption is supported by similar amino acid compositions of microbial biomass across various species [17–19]. By changing the units of concentration of different amino acids, we can make this universal stoichiometry to be given by 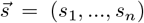. Microbes also uptake essential resources *R*_*i*_ from their environment (black arrows in Fig. 1a), with the uptake rate proportional to *R*_*i*_. Assuming a passive exchange of essential resources between species and the environment, without loss of generality, we can set the maximal uptake rate from the environment as *R*_*i*_, which is the same for each species. Thus, the maximal total flux of the essential resource *i* for species *α* that could in principle be converted to the biomass is given by 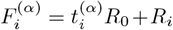. This expression accounts for both the intracellular transformation of the primary resource and the direct uptake of *R*_*i*_ from the environment.

The growth of microbes follows Liebig’s law of the minimum, positing that growth is constrained by a single limiting essential resource [20, 21]: the specific growth rate 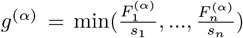. If resource *i* is the limiting resource of species *α*, then for resource *j*, the flux used for biomass synthesis is 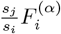 (red arrows in Fig. 1a), and the remaining flux (i.e.,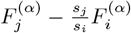) is the overflow flux (orange arrows in Fig. 1a). We then model the population dynamics within a chemostat environment, characterized by constant resource supply rates and fixed dilution rates for both species and resources (Fig. 1b).

### Graphical interpretation of the feasibility and stability criteria

If the transformation efficiency of the primary resource is the same across all species, the transformation vector can be normalized so that 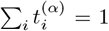. When applying our model to actual data on auxotrophic strains of *E. coli*, we will relax this assumption. In scenarios involving three essential resources, each species can be represented as a point within an equilateral triangle (pink and brown dots on a three-dimensional simplex in Fig. 1c). The distances from the species’ position to the simplex’s three sides (*t*_1_, *t*_2_, and *t*_3_) are proportional to its capacity to transform the primary resource *R*_0_ to essential resources *R*_*i*(*≥*1)_ respectively. Similarly, the stoichiometry vector 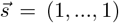 is depicted as the red cross at the center of the equilateral triangle (Fig. 1). These graphical representations, along with subsequent analysis, will be extended to cases where species differ in their transformation efficiencies.

In ecosystems, achieving a stable steady state hinges on two key criteria: feasibility [22] and stability [23, 24]. A state is deemed feasible if all surviving species have zero net growth rates and maintain non-negative abundances at the steady state [22]. In our model, we extend this concept by introducing “flux feasibility” for full-diversity communities. This is determined geometrically by whether the stoichiometry vector 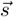 (the red cross in Fig. 2a) lies within the convex hull formed by the species’ transformation vectors 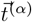 (yellow triangle in Fig. 2a). Intuitively, each species *α* contributes a transformation vector 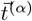, and the community as a whole can adjust species abundances to produce an average transformation flux vector 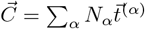, where *N*_*α*_(≥ 0) is the relative abundance of species *α*. If *s* lies outside the convex hull defined by 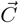, coexistence is not feasible, as no combination of species abundances can realize the required transformation flux (Fig. 2b). Mathematically, a full-diversity community is achievable if 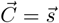. Details of the algebraic analysis for communities with sub-maximal diversity are provided in Supplementary Fig. 1.

**FIG. 2.**
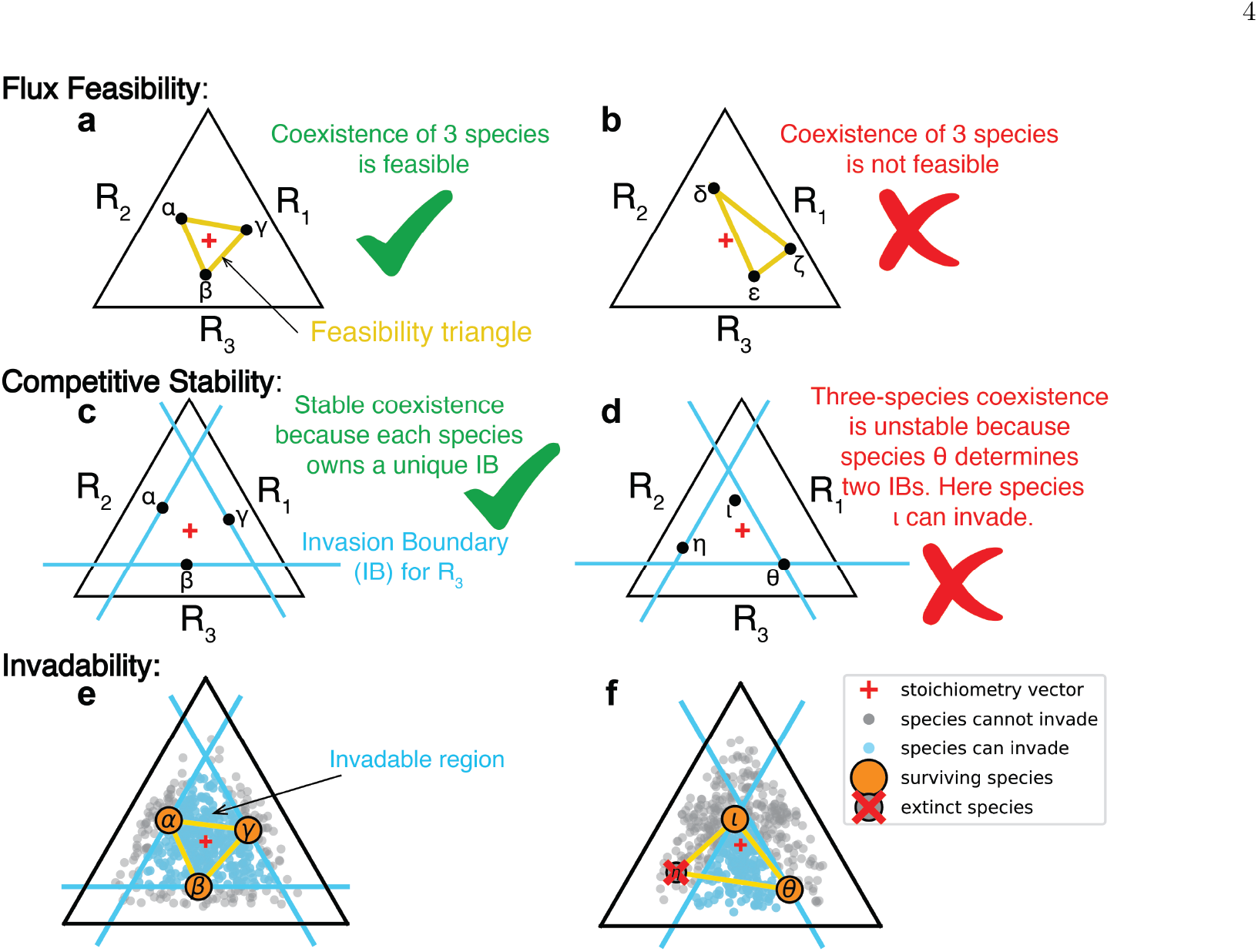
Graphical interpretation of flux feasibility and competitive stability. **a-b** The flux feasibility of a full-diversity community depends on whether the stoichiometry vector is within the convex hull formed by connecting transformation vectors of all species (yellow feasibility triangle). **c-d** The competitive stability of a full-diversity community depends on whether each species is growth-limited by a unique resource. For each essential resource, the species growth-limited by it has the weakest transformation capability for the resource and needs to uptake the resource most from the environment, thereby determining the IB (Invasion Boundary) that is a line drawn through the species' position within a simplex and parallel to the side representing its growth-limiting resource. **e-f** Model simulations demonstrate the invadability. The invadable region is bounded by IBs. These simulations start a community assembly with three species (labeled in each panel) in a chemostat, allowing it to reach a stable steady state. Invasion simulations, conducted 497 times, involve introducing a single species with a randomly chosen transformation vector into the steady state to test its ability to invade the community. Successful and failed invasions are marked with light blue and grey dots, respectively.

In ecological models involving multiple essential resources, a species’ growth rate is typically determined by a single growth-limiting resource [25, 26]. In our model, for a given essential resource *R*_*i*_, the species with the least capacity to produce it on its own must uptake more of it from the environment than other species, thereby becoming growth-limited by this resource. To illustrate this, let us consider a case with three essential resources and three species (Fig. 2c). Geometrically, the species that is growth-limited by a specific resource is the one whose position within the simplex is the closest to the side representing this resource. For instance, in Fig. 2c, species *β* is closest to the triangle’s bottom side, indicating that it is growth-limited by resource *R*_3_. Similarly, species *α* and *γ* are limited by resources *R*_2_ and *R*_1_, respectively. More generally, in a maximally diverse ecosystem with the number of species equal to the number of resources, each species’ growth is limited by a unique resource, and each resource limits the growth of one species only [26]. Here we define “competitive stability” as a state in which: (1) no resource limits the growth of more than one species, and (2) the steady state cannot be invaded by any species that was initially present in the community but extinct at the steady state. This principle holds for the community in Fig. 2c, confirming its “competitive stability”. Conversely, the community shown in Fig. 2d is not competitively stable since the growth of species *θ* is limited by both *R*_1_ and *R*_3_.

This observation allows us to introduce an “Invasion Boundary” (IB), a line (or more generally a hyperplane) drawn through each species’ position within a simplex and parallel to the side representing its growth-limiting resource. Any potential invader species positioned closer to the resource than the IB cannot successfully invade the community. Indeed, for such a species, the concentration of this resource will be insufficient for exponential growth in our chemostat. Conversely, an invader species located further away from all resources than surviving species’ IBs (species within the blue triangle in Fig. 2c) will succeed in invading this community. This theoretical prediction was confirmed by our numerical simulations of ~ 500 attempted invasions of this community by random invader species (successful/failed invasions as blue/grey dots in Fig. 2e).

The scenario in Fig. 2d is inherently unstable, and one can show that species *ι* would eventually outcompete *η* (Supplementary Fig. 1). Two surviving species, *θ* and *ι*, will be growth-limited by *R*_1_ and *R*_2_ correspondingly (see detailed algebraic analysis in Supplementary Fig. 1). As before, we predict that successful invaders have transformation vectors inside the region formed by IBs of surviving species (*θ* and *ι*) in the community, while unsuccessful invaders have transformation vectors outside this region. This theoretical prediction is corroborated by numerical simulations shown in Fig. 2f, where the IBs for the two surviving species separate successful invaders (blue dots) from unsuccessful ones (grey dots) among ~ 500 attempted invasions.

### Model with externally supplied essential resources and different conversion efficiencies

Until now, our analysis has assumed that the primary resource *R*_0_ was the only external supply to the ecosystem. In this case, the only source for essential resources *R*_*i*_ was the transformation of *R*_0_ by species in the community. However, in natural ecosystems, some essential resources may be directly supplied. Our methods for evaluating flux feasibility and competitive stability can be easily extended to this scenario. Externally supplied resources are modeled by a vector, 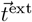, whose *i*’s component is the ratio of the external supply rate of the essential resource *R*_*i*_ to that of the primary resource *R*_0_ (Supplementary Fig. 2a). As a result, the community transformation vector becomes 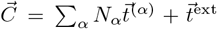, combining the collective transformation of *R*_0_ into *R*_*i*_ with the direct supply of *R*_*i*_. For a maximally diverse community in steady state, the community transformation vector must be equal to the stoichiometry vector 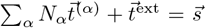. The condition for flux feasibility thus becomes 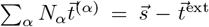 with positive *N*_*α*_. Geometrically, this is depicted as a shift of the stoichiometric vector from 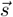 to 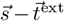 (red star inside a simplex in Supplementary Fig. 2a). The community is feasible provided that this shifted stoichiometry vector 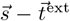 lies within the convex hull formed by 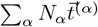, validated by numerical simulations shown in Supplementary Fig. 2. Therefore, the volume of this convex hull can be interpreted as “structural stability” to withstand a wide range of resource supply conditions [27–29].

We also adapted our graphical and algebraic methods to the situation when species do not fully transform the primary resource *R*_0_ to essential resources *R*_*i*_, resulting in different overall conversion efficiencies 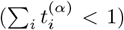. We demonstrated this model variant with a system of 2 essential resources and 2 species (Supplementary Fig. 3). Our graphical and algebraic approach remains applicable in this general case. For the community shown in Supplementary Fig. 3, we predict the coexistence of the two species since they satisfy (1) the flux feasibility condition: each species is positioned on opposite sides of the *y* = *x* line defined by the stoichiometry vector (1,1) (Supplementary Fig. 3a) and (2) the competitive stability condition: each species has a unique limiting resource (Supplementary Fig. 3a). The outcome of our invasion simulations validated the prediction based on our graphical method (invadable region with blue dots in Supplementary Fig. 3c).

### Auxotroph communities are more resilient to resource fluctuations

Why is auxotrophy so widespread in natural communities? What advantage could there be for a prototrophic species, capable of synthesizing all essential resources for its growth, to lose this ability to produce one or more resources? Traditional explanations typically invoke evolutionary arguments, such as the Black Queen Hypothesis or genome reduction, which suggest that organisms may lose costly biosynthetic functions when those functions are shared by others in the community [30, 31]. In contrast, our model offers an alternative, ecological answer: auxotrophy can expand the feasibility volume– the range of resource supply conditions that support stable coexistence–thus promoting community-level structural stability. By comparing the relatively small feasibility volume of communities populated by prototrophic species (small yellow triangle in Fig. 3a) to the progressively larger feasibility volumes of communities composed of single-resource auxotrophs (medium yellow triangle in Fig. 3c) and double-resource auxotrophs (largest possible yellow triangle in Fig. 3e), we conclude that auxotrophy may enhance community survival in environments with fluctuating external supplies of essential resources. Our graphical approach illustrates that changes in the external supply of essential resources manifest as shifts in the vector 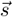. Communities with a larger feasibility domain are better equipped to handle these shifts, as all community species can survive as long as the shifted vector 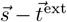 remains within the feasibility domain.

**FIG. 3.**
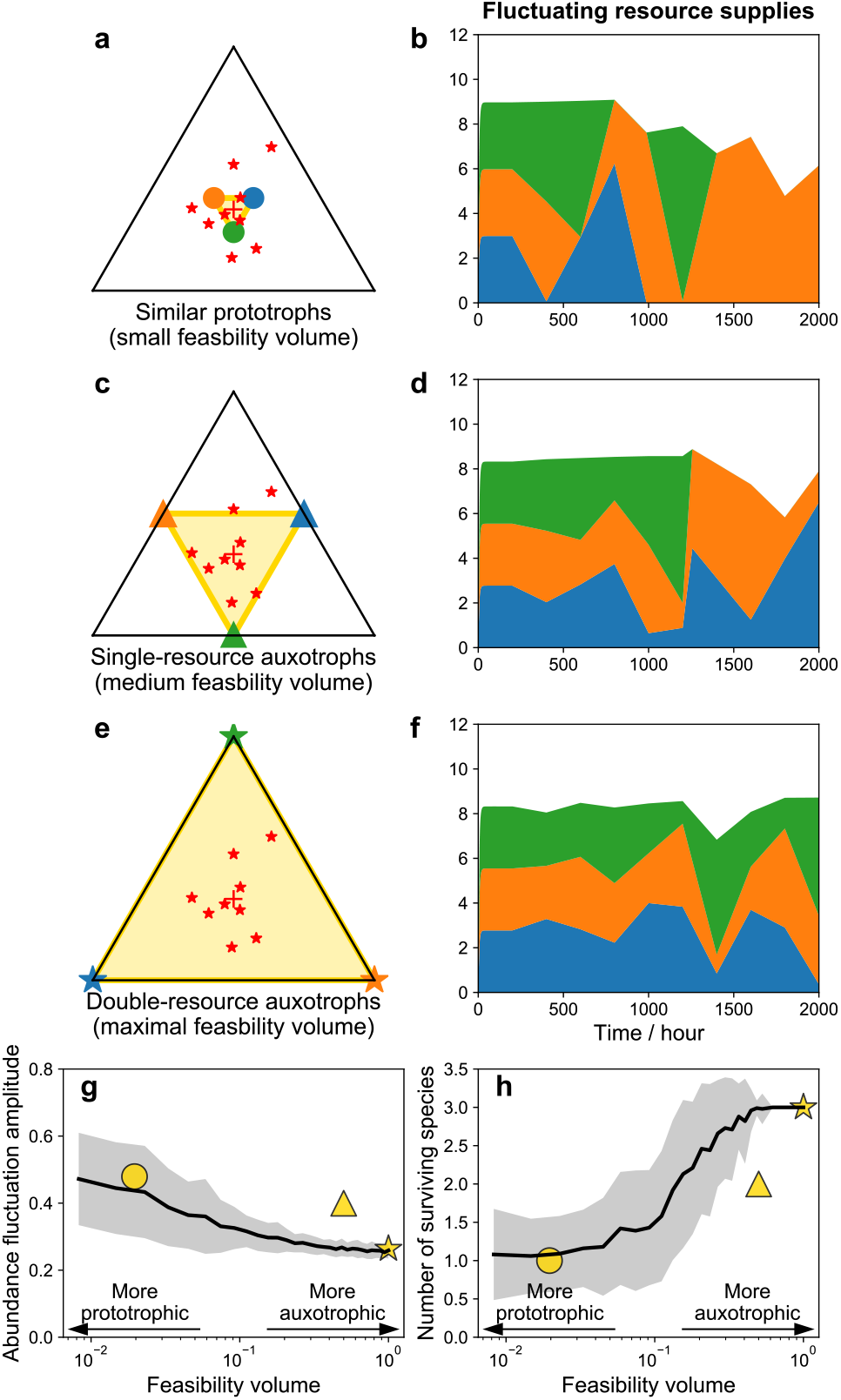
Auxotroph communities are more resilient to resource fluctuations. In these simulations, fluxes of essential resources externally supplied to the system change every 200 hours, with each change gradually increasing in magnitude. The red stars represent 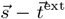 after each such change. **a** A community of three prototrophs with similar transformation abilities. **b** The dynamics of the community composition shown in panel a under fluctuating supply of essential resources. **c** A community of three single-resource auxotrophs. **d** The compositional dynamics of the community shown in panel c. **e** A community of three double-resource auxotrophs. **f** The compositional dynamics of the community shown in panel e. **g** The average magnitude of fluctuations in microbial abundances, defined as the standard deviation of time-series abundances divided by their mean, decreases as the feasibility volume increases from prototrophs to double-resource auxotrophs. **h** The average richness of assembled communities increases with the feasibility volume. Simulations in panels e and f are averaged over 100 repeats for each bin of feasibility volumes. The black line represents the mean values of the y-axis and the grey area represents one standard deviation from the mean. Simulations shown in panels a-f are indicated by the corresponding symbols: circle for prototrophs in panel a, triangle for single-resource auxotrophs in panel c, and star for double resource-auxotrophs in panel e.

To verify this intuition, we studied the dynamics of three communities under conditions of fluctuating essential resource supplies, each with varying feasibility volumes: (1) communities with similar prototrophic species (Figs. 3a-b), (2) communities of single-resource auxotrophs with medium feasibility volume (Figs. 3c-d), and (3) communities comprising double-resource auxotrophs with maximal feasibility volume (Figs. 3e-f). We modeled these fluctuations by independently and uniformly sampling the ratio of the supply rate of each essential resource to that of the primary resource 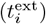 between 0 and 0.3 every 200 hours.

When three prototrophic species with similar transformation abilities face fluctuations in essential resource supplies (Fig. 3a-b), their species abundances fluctuate dramatically. In a representative simulation shown in Fig. 3b, the blue and green species die out after 1400 hours due to resource fluctuations. In contrast, in a community of three single-resource auxotrophs (Fig. 3c-d), three species manage to coexist longer, with two surviving even after 2000 hours. The community composed of three double-resource auxotrophs has the largest possible feasibility volume, so in a representative simulation of this community shown in Fig. 3f, all three species survived indefinitely. We systematically explored how the feasibility volume impacts the abundance fluctuation and the richness of the assembled community. By measuring ‘abundance fluctuation amplitude’ through the ratio of the standard deviation to the mean of species abundances, we observed a consistent decrease in abundance fluctuation amplitude with increasing feasibility volume (Fig. 3g). This decrease reduces the likelihood of species extinction, thereby enhancing community richness and stability (Fig. 3h).

### Accurate predictions of experimentally assembled auxotroph communities

To showcase our model’s capability in predicting community assembly outcomes, we applied it to experimental data from a synthetic community comprised of 14 *E. coli* auxotroph strains, each genetically engineered to lack the ability to synthesize one specific amino acid [10]. This synthetic community was investigated through (1) pairwise co-cultures of auxotroph pairs (91 pairs in total, 1 replicate each pair) and (2) a co-culture of all 14 auxotrophs (2 replicates). We first fitted the transformation vectors for these auxotrophs using fold-growth observed in pairwise co-culture growth experiments (Supplementary Fig. 5). Using these fitted vectors, we then predicted the assembly outcome of the community comprising all auxotrophs. Instead of fitting the stoichiometry vector, we used the previously reported biomass composition of amino acids in *E. coli* [10].

We iteratively adjusted the transformation vectors for each of the 14 strains across every amino acid (14 *×* 13 = 182 parameters since each synthetic auxotroph strain is unable to synthesize one amino acid, which results in its transformation value of zero for that amino acid). The fitting used the data on fold-growth in 91 pairwise co-culture experiments (Supplementary Fig. 5; see Methods section for details). Note that we use the version of our model that allows for the incomplete transformation of the primary resource to essential resources 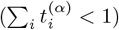, which can be caused by the leakage of essential resources [32] and the leakage of essential resources [33, 34]. To illustrate our iterative algorithm, let us consider a co-culture of Δ*T* and Δ*M* strains. If the fold-growth of Δ*T* predicted by our model exceeds or falls short of the experimental value, we adjust Δ*M* ‘s transformation ability of *T* upward or downward respectively to minimize the discrepancy (Supplementary Fig. 4). This process is repeated until we obtain the best-fitted transformation vectors (Fig. 4b and Supplementary Fig. 6).

**FIG. 4.**
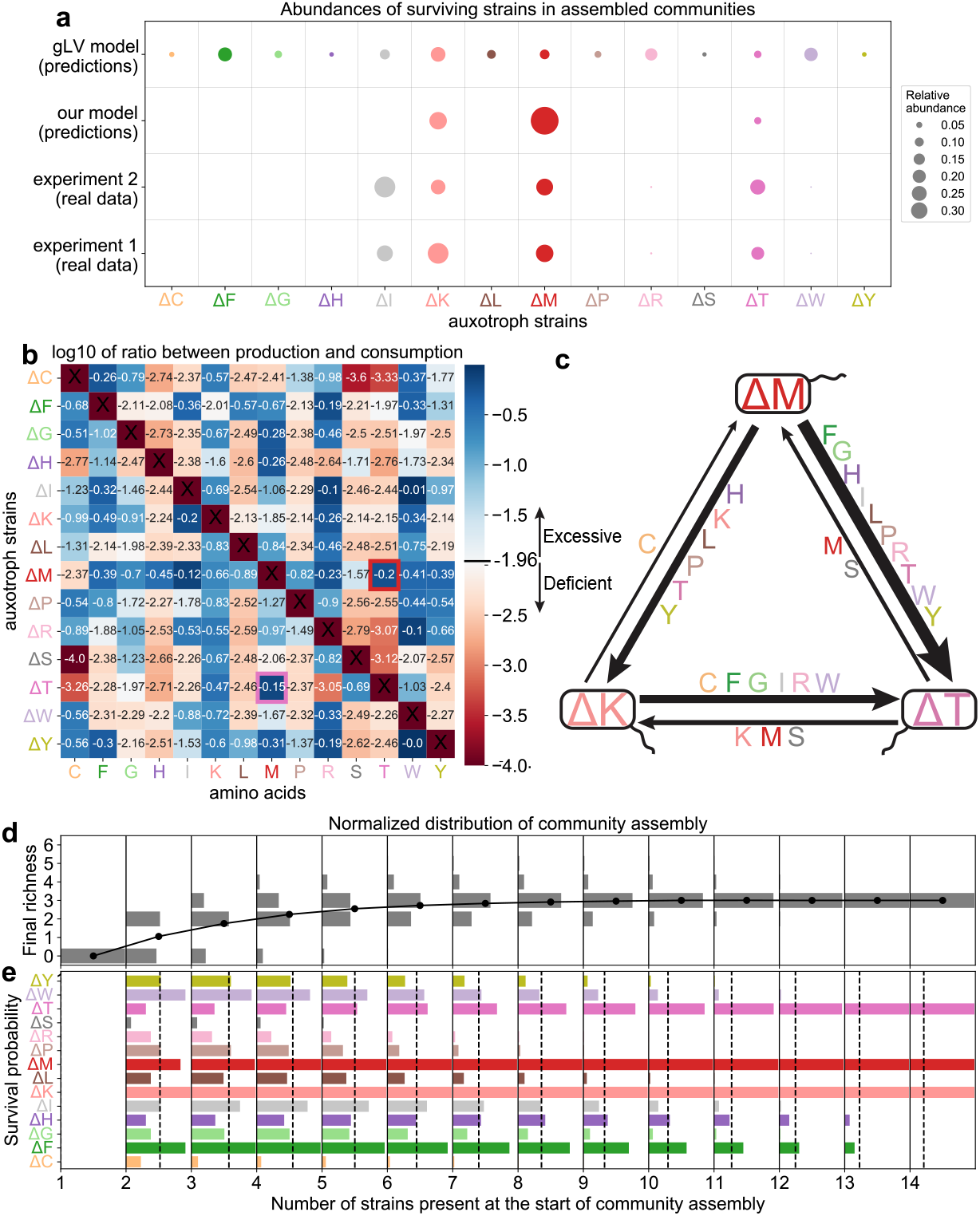
Our model prediction of the community assembly inoculated by *E. coli* synthetic auxotrophs. Color assignments for all strains and amino acids are consistent across all panels. The full names of amino acids are listed in the Methods section. The prefix Δ indicates a single amino acid auxotrophy (e.g., Δ*T* refers to the strain unable to synthesize Threonine (T)). **a** Our model's prediction of the steady state composition for the full diversity 14-member assembly (second line) closely matches the experimental results (two biological replicates shown in lines 3 and 4) by correctly predicting three out of four surviving strains. Conversely, the generalized Lotka-Volterra (gLV) model fitted to the pairwise assembly data (first line) incorrectly predicts the survival of all 14 strains. **b** The ratio of the best-fitted transformation vector in our model to the stoichiometry vector experimentally measured in *E. coli*. The synthetic resource auxotrophy for each strain is marked with ×. Red and purple squares mark the only pair of strains (Δ*M* and Δ*T*) with a strong (*>* 0.5 for each strain) mutual exchange flux of amino acids (T and M) they cannot produce. **c** The exchange fluxes of amino acids between three surviving strains predicted by our model. **d** Distribution of the final community richness when different subsets of *E. coli* strains are initialized with equal amounts to start the community assembly. A separate distribution is shown for each initial richness. Dots connected by the black line represent the average final richness for each initial richness. **e** For each initial richness, the survival probability of each strain is computed by measuring the fraction of combinations in which it survives. The black dashed line shows the average survival probability across all strains.

Using these fitted transformation vectors, our model accurately predicted three strains out of four surviving strains observed experimentally (Δ*K*, Δ*M*, and Δ*T* in Fig. 4a) and the extinction of the remaining 10 strains. For comparison, we applied the previously proposed gLV model [10], which has the same number of fitting parameters (182) as our model and used the same pairwise co-culture data for fitting (see Methods for more details). Although the gLV model fitted pairwise co-culture data better (Supplementary Fig. 5), it completely failed in selecting which strain would survive in the full community assembly. Indeed, the gLV model predicted the survival for all 14 strains, diverging significantly from experimental observations with only four survivors (gLV results in Fig. 4a). This spectacular failure of the gLV model, which only accounts for pairwise interactions, indicates the importance of higher-order interactions in auxotroph communities. We also simulated our model in batch cultures with a dilution ratio of 100 (more details in the Methods section), which better reflects the experimental setup [10]. We found the same three strains (Δ*K*, Δ*M*, and Δ*T*) survive with their respective relative abundances of 0.246, 0.718, and 0.036, closely matching the result of our chemostat model (0.242, 0.700, and 0.058). The fitted transformation vectors enable us to accurately map the niche of each auxotroph within the resource space, identify strong competitors, and identify the most effective mutualistic pairs or trios (Fig. 4b).

Notably, the strain Δ*T* emerges as the ideal mutualistic partner for the strain Δ*M*, due to the surplus production of *M* by Δ*T* (the purple square in Fig. 4b) and abundant production of *T* by Δ*M* (the red square in Fig. 4b). We quantified the cooperative potential between two strains, Δ*X* and Δ*Y*, by the product 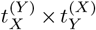, which captures their mutual reliance on each other’s amino acid production (Supplementary Fig. 7). Across all pairs, Δ*T* and Δ*M* display the strongest cooperation potential, with 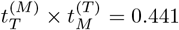, the highest value observed (Supplementary Fig. 7). We further summarized the exchange of amino acids between surviving strains using a graph (Fig. 4c), revealing that Δ*M* is a key supplier of most amino acids to both Δ*K* and Δ*T*. Meanwhile, Δ*K* provides two other strains with cysteine (C), which neither of them can synthesize in adequate amounts. While our model accurately recapitulates the survival of most strains in the 14-member auxotroph community, it does not predict the survival of strain Δ*I*, which was observed experimentally. We checked the limiting amino acids when Δ*K*, Δ*M*, and Δ*T* coexist, finding that the limiting amino acids are C, M, and S. Δ*I* cannot survive due to its ability to produce S is weaker than Δ*K*, Δ*M*, and Δ*T*. Indeed, the production-consumption ratio of the amino acid S for the strain Δ*I* (0.003) is smaller than 0.007 for Δ*K*, 0.027 for Δ*M*, and 0.204 for Δ*T* (Fig. 4b). Under real experimental conditions, Δ*I* might survive because of factors our model does not include, like the availability of other resources or environmental details not captured here.

We also simulated community assembly from every possible subset of the 14 *E. coli* synthetic auxotroph strains (Fig. 4d-e). We found that the mean richness in the finally assembled communities increases with the number of strains present initially (Fig. 4d). As the initial richness increases, Δ*K*, Δ*M*, and Δ*T* are more frequently assembled (Fig. 4e). As a null expectation, we simulated 10,000 assemblies of 14 single-resource auxotroph strains with randomly assigned transformation vectors. These random assemblies consistently achieved higher final richness than the assembly result of 14 *E. coli* strains, underscoring the uniqueness of these *E. coli* strains (Supplementary Fig. 8).

## Discussion

The role of auxotrophic species in shaping ecological dynamics and community assembly has been underexplored in mechanistic models. In this study, we introduced an innovative ecological model that incorporates both the catabolic and anabolic processes of auxotrophic and prototrophic species, as well as their interactions with the environment. This model provides both graphical and algebraic interpretations of flux feasibility, emphasizing the community’s strategies to optimize species abundances to minimize essential resource overflow. It elucidates why auxotroph communities are more diverse and resilient to resource fluctuations. For competitive stability in such optimized communities, species must occupy distinct niches characterized by non-overlapping limiting resources. Applied to experimental data from a synthetic community of *E. coli* auxotroph strains, our model successfully predicted the survival of three out of four strains observed.

Previous attempts to quantitatively understand interactions between auxotrophs have primarily utilized gLV models [10]. However, these models completely failed to predict the assembly of a diverse 14-strain community. This failure arises because gLV models fitted to pairwise data capture only cooperative interactions, due to the mutually beneficial exchange of missing amino acids. Consequently, all interaction coefficients inferred from such models are positive. When such a model is applied to describe a full-diversity community, it erroneously predicts the survival of all strains. A more realistic approach necessitates incorporating both negative, competitive interactions and accounting for complex higher-order effects. Our model addresses these shortcomings by introducing concepts such as competitive stability, invasion boundaries, and growth-limiting resources. These elements not only capture the competition between species but also highlight the potential for one species to drive another to extinction, thereby offering a more nuanced and realistic view of species interactions and community dynamics.

Traditional consumer-resource models [15, 25, 26, 35] have another deficiency: they usually neglect the details of intracellular metabolism. In contrast, our model offers a coarse-grained yet realistic description of intracellular metabolic transformations, transforming a primary resource (e.g., glucose) into multiple metabolites essential for biomass production (e.g., amino acids), akin to flux balance analysis (FBA) [36, 37]. However, our model diverges from FBA by not focusing on maximizing each species’ growth rate. Instead, it emphasizes the cooperative sharing of essential resources and balances their overall production and consumption fluxes only at the community level (i.e., flux feasibility). Our model also accounts for the competition between species for niches defined by growth-limiting resources.

Similar to our previous intermediate-scale model [16], here we use a simplified CRM framework that focuses on resource transformation and acquisition mechanisms, while ignoring previously considered details such as growth inhibition from resource accumulation and constant leakage of resources. This simplification makes graphical and algebraic analyses tractable, which were not feasible before. Moreover, the simplified model dramatically reduces the number of parameters, enabling the prediction of 14-member community assembly outcomes based solely on pairwise co-culture data.

Additionally, our model provides insights into the prevalence of auxotrophs in natural communities [1–4], clarifying their role in enhancing community stability [5]. By relying on external sources for essential nutrients, auxotrophs foster interdependence among species, serving as a stabilizing force within the community. We model this interdependence by matching the excess production with the growth limitations of the surviving species. When the external supply of resources shifts, auxotrophic species in our model adjust their populations to buffer these fluctuations, thereby mitigating the impact of changes in resource availability. Thus, communities rich in auxotrophs often display greater ecological resilience and stability compared to those dominated by prototrophs.

Our model demonstrates that an auxotrophic community as a whole needs to adjust species abundances so that its community transformation vector 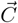 best aligns with the biomass stoichiometry 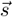. As a result, community composition and growth depend on the combined supply of several essential resources rather than any single one. This parallels modern co-limitation theory, which posits that microbial growth could be jointly constrained by multiple resources [38, 39]. Fully exploring this link between auxotroph ecology and quantitative co-limitation falls beyond our present scope and remains a promising direction for future work.

In this work, we assumed identical stoichiometries for biomass synthesis across all species. While this is a reasonable approximation for communities composed of closely related species, it can be a limitation when modeling more phylogenetically diverse communities. To evaluate the impact of unequal stoichiometries, we simulated community assembly using different subsets of 14 single-resource auxotrophic species, each assigned a stoichiometry vector derived by multiplying *E. coli* ‘s stoichiometry by a random number drawn from a uniform distribution between 0.5 and 1.5. We compared two scenarios: (1) all species had the same stoichiometry, and (2) species had randomly varied stoichiometries. For both cases, we varied the number of species present at the start of the community assembly (i.e., the initial richness) and recorded the final richness after the community assembly. We observed no big difference in the final richness between the same and varied stoichiometries (Supplementary Fig. 9a,c). It is important to note that our current graphical and algebraic analyses rely on the assumption of equal stoichiometry and cannot be directly applied when stoichiometries vary. Moreover, previous work has demonstrated that when stoichiometries differ, consumer-resource models with multiple essential resources can exhibit multi-stability and regime shifts [25, 40]. We also compared the difference in the final assembled community richness between equal and unequal uptake rates and did not observe a difference (Supplementary Fig. 9a,b). Additionally, our model assumes that overflow resources are solely used for biomass production and does not account for their potential catabolism as energy, carbon, or nitrogen sources. Including these aspects represents an important direction for future work.

## METHODS

### Datasets

The experimental pairwise and 14-member cocultures of *E. coli* auxotroph strains can be found in the Supplemental Information of Mee et al [10]. The abbreviations of amino acids are as follows: Arginine (R), Cysteine (C), Glycine (G), Histidine (H), Isoleucine (I), Leucine (L), Lysine (K), Methionine (M), Phenylalanine (F), Proline (P), Serine (S), Threonine (T), Tryptophan (W), and Tyrosine (Y).

### The ecological model with production and overflow of essential resources

Our model considers the depletion, transformation, uptake, and overflow of resources as well as the growth of microbes. For microbial species *α*, it consumes the sole primary resource *R*_0_ and then transforms it into essential resources *R*_*i*(*≥*1)_. As a result, the transformation flux for essential resource *i* by species *α* is 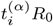, where *R*_0_ is the concentration of the primary resource and 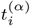 is the rate at which the primary resource *R*_0_ is transformed to essential resource *i*. Additionally, species *α* can uptake resource *i* directly from the environment, with its uptake flux proportional to *R*_*i*_. Assuming that microbial species passively uptake all essential resources at the same rate, we set the uptake resource as *R*_*i*_. The total influx of essential resource *i* is given by 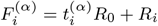.

The specific growth rate of species *α* is described by Liebig’s law of the minimum: 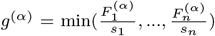 where *s*_*i*_ is the stoichiometric fraction of essential resource *i* required for the biomass synthesis. If the influx of resource *i* for species *α* exceeds its rate consumed for biomass synthesis (i.e.,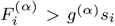), the unconsumed part forms the overflow flux (i.e.,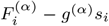). The dynamics of species abundances (denoted as *N*_*α*_) and resource concentrations are described by the following equations:

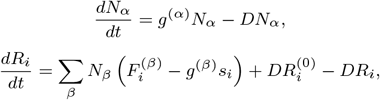

where *D* is the dilution rate and 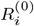 is the pool concentration of resource *i*.

For simulations involving fluctuations in essential resources, the ratio 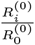 for each essential resource *i >* 0 is randomly sampled from a uniform distribution between 0 and 0.3 every 200 hours.

Regarding the data with 14 *E. coli* auxotroph strains [10], fold-growth observed in pairwise co-cultures is used to fit 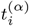. Specifically, we used the gradient descent algorithm to minimize the sum of the squared differences between the log10-transformed fold-growths predicted by the model and those observed experimentally. For each pairwise co-culture of Δ*X* and Δ*Y*, if the model overestimates or underestimates Δ*X*’s fold-growth, we adjust Δ*Y* ‘s transformation ability of X downward or upward accordingly to reduce the fitting error. All simulations of single-batch pairwise co-cultures start with two strains with equal abundances, consistent with the experimental setup [10]. Then our model was run with the dilution rate *D* = 0 over 84 hours to obtain model-predicted fold-growths. *D* = 0 was used in our chemostat model to simulate the batch growth. Eventually, the fitted parameters are used to predict the assembly outcome for the 14-member co-culture. Similar to the experimental setup [10], we initialized the simulation of the 14-member co-culture with equal amounts for all strains. Beyond the chemostat we laid out, we also simulated the case of serial dilution: analogous to as the experimental 14-member coculture [10], we simulated this case daily using our model by (1) assuming dilution rate *D* = 0 and (2) stopping the simulation every 24 hours, diluting microbial abundances and resource concentrations by a factor of 100, and replenishing with the primary resource.

### The gLV model and its fitting

The gLV model we utilized is from the mathematical model proposed in Mee et al [10]. In this model, the abundance dynamics of strain *α* is specified as

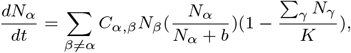

where *N*_*α*_ is the abundance of strain *α, C*_*α*,*β*_ is the cooperativity coefficient, *K* is the carrying capacity which is assumed to be 10^9^, and *b* is the buffer term used for low-density growth which is assumed to be 10^5^. Fold-growth in pairwise co-cultures is used to fit *C*_*α*,*β*_, whose values are adjusted based on the gradient descent approach to minimize the difference between experimental data and model predictions. Eventually, the model with fitted *C*_*α*,*β*_ is used to generate the prediction for the 14-member co-culture result.

### Statistics

All simulations and analyses were performed using standard numerical and scientific computing libraries in the Python programming language (version 3.7.1) and Jupyter Notebook (version 6.1).

### Data and code availability

All code for simulations used in this manuscript can be found at https://github.com/wt1005203/auxotroph.

## Acknowledgements

We thank Veronika Dunbinka and Zihan Wang for useful discussion.

## Author contributions

All authors helped devise the study; TW performed the research and analyzed the data; SM supervised the research; TW and SM wrote the manuscript.

## Competing Interests

The authors declare no competing interests.

## Notes

### Competing Interest Statement

The authors have declared no competing interest.

### Summary of Updates

Figure 4 revised; author affiliations updated; new simulation results added to the Supplemental Information.

### Funding

- This research was supported by startup funds from the Department of Biological Sciences at Purdue University (to T.W.). This research was also supported in part by grants from the NSF (DMS-2235451) and Simons Foundation (MPS-NITMB-00005320) to the NSF-Simons National Institute for Theory and Mathematics in Biology (NITMB).

## References

[1] G. D’Souza, S. Waschina, S. Pande, K. Bohl, C. Kaleta, and C. Kost, Less is more: selective advantages can explain the prevalent loss of biosynthetic genes in bacteria, Evolution 68, 2559 (2014).

[2] M. Embree, J. K. Liu, M. M. Al-Bassam, and K. Zengler, Networks of energetic and metabolic interactions define dynamics in microbial communities, Proceedings of the National Academy of Sciences 112, 15450 (2015).

[3] Y.-F. Liu, D. D. Galzerani, S. M. Mbadinga, L. S. Zaramela, J.-D. Gu, B.-Z. Mu, and K. Zengler, Metabolic capability and in situ activity of microorganisms in an oil reservoir, Microbiome 6, 1 (2018).

[4] W. R. Harcombe, J. M. Chacón, E. M. Adamowicz, L. M. Chubiz, and C. J. Marx, Evolution of bidirectional costly mutualism from byproduct consumption, Proceedings of the National Academy of Sciences 115, 12000 (2018).

[5] S. Starke, D. M. Harris, J. Zimmermann, S. Schuchardt, M. Oumari, D. Frank, C. Bang, P. Rosenstiel, S. Schreiber, N. Frey, et al., Amino acid auxotrophies in human gut bacteria are linked to higher microbiome diversity and long-term stability, The ISME Journal 17, 2370 (2023).

[6] L. Oña, S. Giri, N. Avermann, M. Kreienbaum, K. Thormann, and C. Kost, Obligate cross-feeding expands the metabolic niche of bacteria. nat ecol evol 5: 1224–1232 (2021).

[7] D. Dutta and S. Saini, Cell growth model with stochastic gene expression helps understand the growth advantage of metabolic exchange and auxotrophy, Msystems 6, 10 (2021).

[8] J. A. Metz, S. A. Geritz, G. Meszéna, F. J. Jacobs, and J. S. Van Heerwaarden, Adaptive dynamics: a geometrical study of the consequences of nearly faithful reproduction, (1995).

[9] B. Momeni, L. Xie, and W. Shou, Lotka-volterra pairwise modeling fails to capture diverse pairwise microbial interactions, Elife 6, e25051 (2017).

[10] M. T. Mee, J. J. Collins, G. M. Church, and H. H. Wang, Syntrophic exchange in synthetic microbial communities, Proceedings of the National Academy of Sciences 111, E2149 (2014).

[11] R. MacArthur, Species packing and competitive equilibrium for many species, Theoretical population biology 1, 1 (1970).

[12] R. Marsland III, W. Cui, and P. Mehta, The minimum environmental perturbation principle: A new perspective on niche theory, The American Naturalist 196, 291 (2020).

[13] M. Raatz, Provision of essential resources as a persistence strategy in food webs, Peer Community Journal 3 (2023).

[14] M. N. Guillen, B. Rosener, S. Sayin, and A. Mitchell, Assembling stable syntrophic escherichia coli communities by comprehensively identifying beneficiaries of secreted goods, Cell Systems 12, 1064 (2021).

[15] R. Marsland III, W. Cui, J. Goldford, A. Sanchez, K. Korolev, and P. Mehta, Available energy fluxes drive a transition in the diversity, stability, and functional structure of microbial communities, PLoS computational biology 15, e1006793 (2019).

[16] C. Liao, T. Wang, S. Maslov, and J. B. Xavier, Modeling microbial cross-feeding at intermediate scale portrays community dynamics and species coexistence, PLoS computational biology 16, e1008135 (2020).

[17] T. Okayasu, M. Ikeda, K. Akimoto, and K. Sorimachi, The amino acid composition of mammalian and bacterial cells, Amino Acids 13, 379 (1997).

[18] H. Akashi and T. Gojobori, Metabolic efficiency and amino acid composition in the proteomes of escherichia coli and bacillus subtilis, Proceedings of the National Academy of Sciences 99, 3695 (2002).

[19] E. M. Heizer Jr, D. W. Raiford, M. L. Raymer, T. E. Doom, R. V. Miller, and D. E. Krane, Amino acid cost and codon-usage biases in 6 prokaryotic genomes: a whole-genome analysis, Molecular biology and evolution 23, 1670 (2006).

[20] C. You, H. Okano, S. Hui, Z. Zhang, M. Kim, C. W. Gunderson, Y.-P. Wang, P. Lenz, D. Yan, and T. Hwa, Coordination of bacterial proteome with metabolism by cyclic amp signalling, Nature 500, 301 (2013).

[21] M. Zampieri, M. Horl, F. Hotz, N. F. Müller, and U. Sauer, Regulatory mechanisms underlying coordination of amino acid and glucose catabolism in escherichia coli, Nature communications 10, 1 (2019).

[22] M. Dougoud, L. Vinckenbosch, R. P. Rohr, L.-F. Bersier, and C. Mazza, The feasibility of equilibria in large ecosystems: A primary but neglected concept in the complexity-stability debate, PLoS computational biology 14, e1005988 (2018).

[23] R. M. May, Will a large complex system be stable?, Nature 238, 413 (1972).

[24] S. Allesina and S. Tang, Stability criteria for complex ecosystems, Nature 483, 205 (2012).

[25] V. Dubinkina, Y. Fridman, P. P. Pandey, and S. Maslov, Multistability and regime shifts in microbial communities explained by competition for essential nutrients, Elife 8, e49720 (2019).

[26] D. Tilman, Resource competition and community structure, 17 (Princeton university press, 1982).

[27] R. P. Rohr, S. Saavedra, and J. Bascompte, On the structural stability of mutualistic systems, science 345, 1253497 (2014).

[28] A. Aparicio, T. Wang, S. Saavedra, and Y.-Y. Liu, Feasibility in macarthur’s consumer-resource model, Theoretical Ecology 16, 225 (2023).

[29] Z. Wang, Y. Fu, A. Goyal, and S. Maslov, Fitness advantage of sequential metabolic strategies emerges from community interactions in strongly fluctuating environments, bioRxiv, 2024 (2024).

[30] J. J. Morris, R. E. Lenski, and E. R. Zinser, The black queen hypothesis: evolution of dependencies through adaptive gene loss, MBio 3, 10 (2012).

[31] S. J. Giovannoni, J. Cameron Thrash, and B. Temperton, Implications of streamlining theory for microbial ecology, The ISME journal 8, 1553 (2014).

[32] K. Shimizu and Y. Matsuoka, Regulation of glycolytic flux and overflow metabolism depending on the source of energy generation for energy demand, Biotechnology Advances 37, 284 (2019).

[33] N. Paczia, A. Nilgen, T. Lehmann, J. Gätgens, W. Wiechert, and S. Noack, Extensive exometabolome analysis reveals extended overflow metabolism in various microorganisms, Microbial cell factories 11, 1 (2012).

[34] J. F. Yamagishi, N. Saito, and K. Kaneko, Advantage of leakage of essential metabolites for cells, Physical Review Letters 124, 048101 (2020).

[35] A. Posfai, T. Taillefumier, and N. S. Wingreen, Metabolic trade-offs promote diversity in a model ecosystem, Physical review letters 118, 028103 (2017).

[36] K. J. Kauffman, P. Prakash, and J. S. Edwards, Advances in flux balance analysis, Current opinion in biotechnology 14, 491 (2003).

[37] J. D. Orth, I. Thiele, and B. Ø. Palsson, What is flux balance analysis?, Nature biotechnology 28, 245 (2010).

[38] W. S. Harpole, J. T. Ngai, E. E. Cleland, E. W. Seabloom, E. T. Borer, M. E. Bracken, J. J. Elser, D. S. Gruner, H. Hillebrand, J. B. Shurin, et al., Nutrient colimitation of primary producer communities, Ecology letters 14, 852 (2011).

[39] N. A. Held, A. Krishna, D. Crippa, R. R. Battaje, A. J. Devaux, A. Dragan, and M. Manhart, Nutrient colimitation is a quantitative, dynamic property of microbial populations, Proceedings of the National Academy of Sciences 121, e2400304121 (2024).

[40] J. Huisman and F. J. Weissing, Biodiversity of plankton by species oscillations and chaos, Nature 402, 407 (1999).

