## Supplemental Information for "Higher-Order Interactions in Auxotroph Communities Enhance Their Resilience to Resource Fluctuations"

### Supplementary Figures

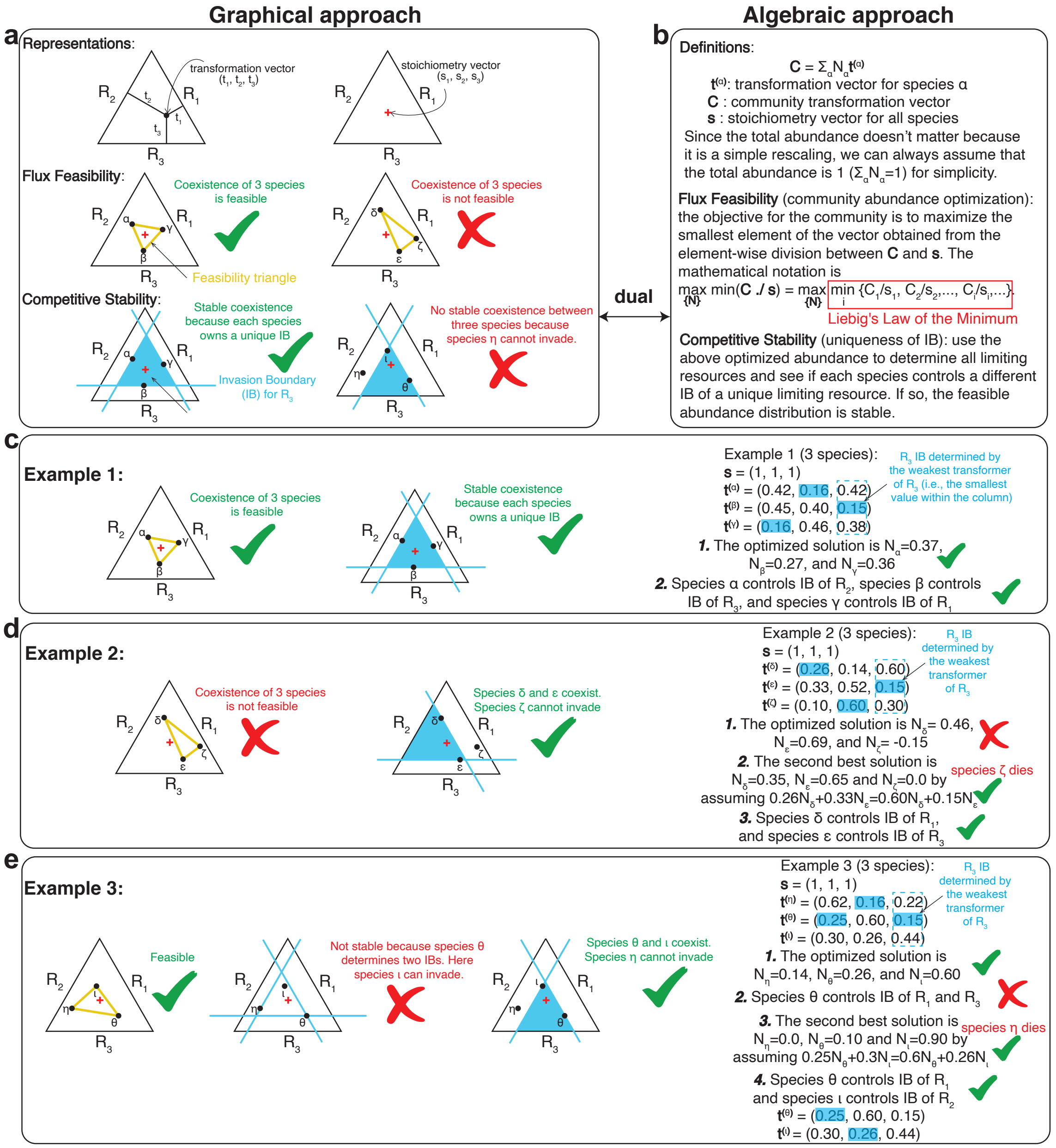


Supplementary Figure 1. **The interpretation of flux feasibility and competitive stability either graphically or through linear algebra. a** When all species are assumed to have the same stoichiometry vector, species' living strategies are determined by their transformation vectors. When $\sum_i t_i^{(\alpha)} = 1$ for all species $\alpha$, each species can be visualized as a dot on the simplex. Positions of all species would help us to generate concepts such as feasibility triangle (yellow triangle) and IBs (Invading Boundaries; blue lines) and further use those concepts to determine flux feasibility and competitive stability. **b** The graphical intuition can be rigorously written using linear algebra where the feasibility criterion is the community abundance optimization and the stability criterion is the uniqueness of IBs for all species. **c-e** Three examples to demonstrate the graphical and algebraic approaches. Point 2 in panel d and point 3 in panel illustrate explicit feasible solutions to the community abundance optimization problem (i.e., $max_{\{\vec{N}\}} min(\vec{C}./\vec{s})$ in panel b). For instance, the expression $0.26N_\delta + 0.33N_\epsilon = 0.60N_\delta + 0.15N_\epsilon$ represents a specific linear combination of species abundances that yields a community transformation vector $\vec{C}$ most closely aligned with $\vec{s}$. These coefficients shown are determined based on the transformation vectors of the species included in the community.

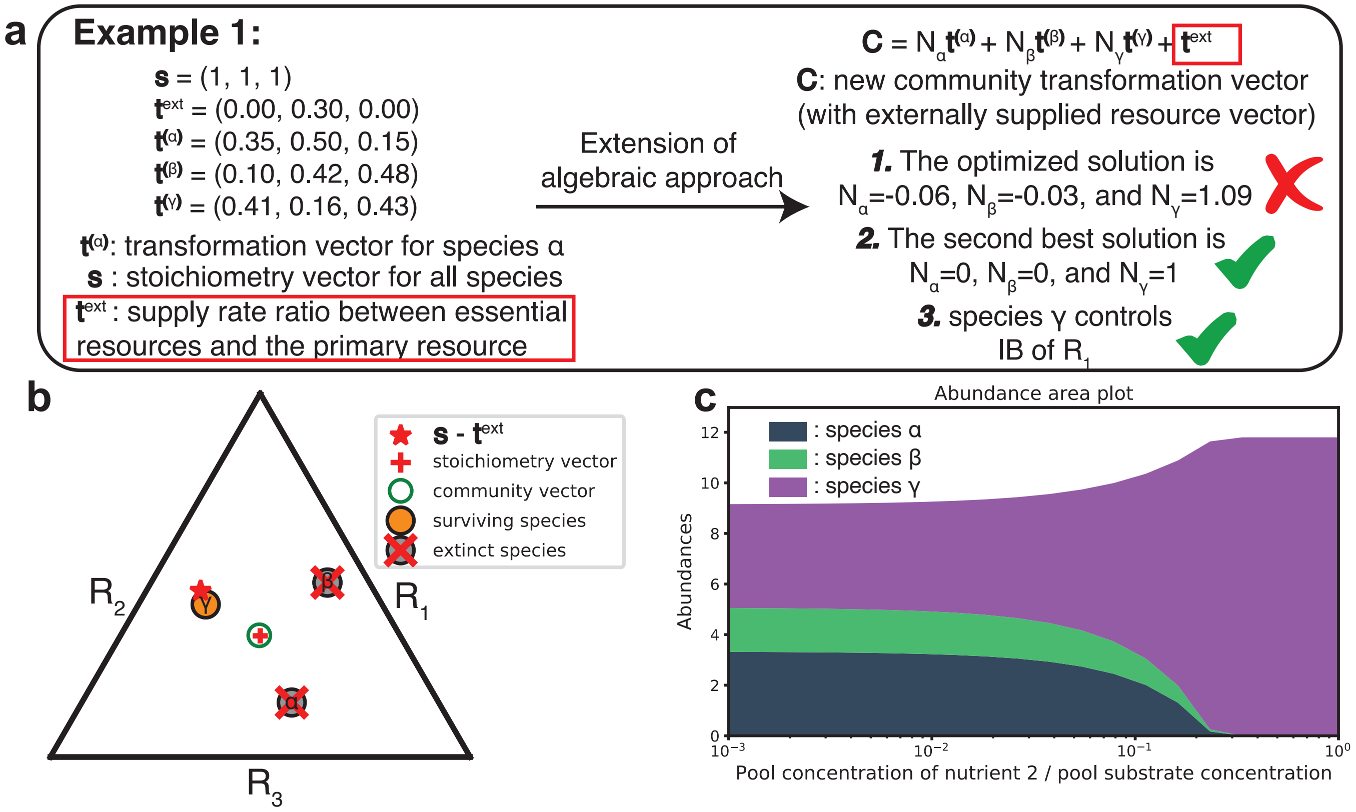


Supplementary Figure 2. **The influence of externally supplied essential resources and the extension of algebraic approach.** Some essential resources can be externally supplied directly into the chemostat. In this example, only resource 2 is externally supplied. **a** The extended algebraic approach can find the stable steady state. **b** The model simulation confirms the prediction for the sample in panel a. The new community vector that includes the contribution of external resource supplies is optimized. **c** The abundance distribution is influenced by the external supply concentration of resource 2. As the supply concentration increases, the diversity drops.

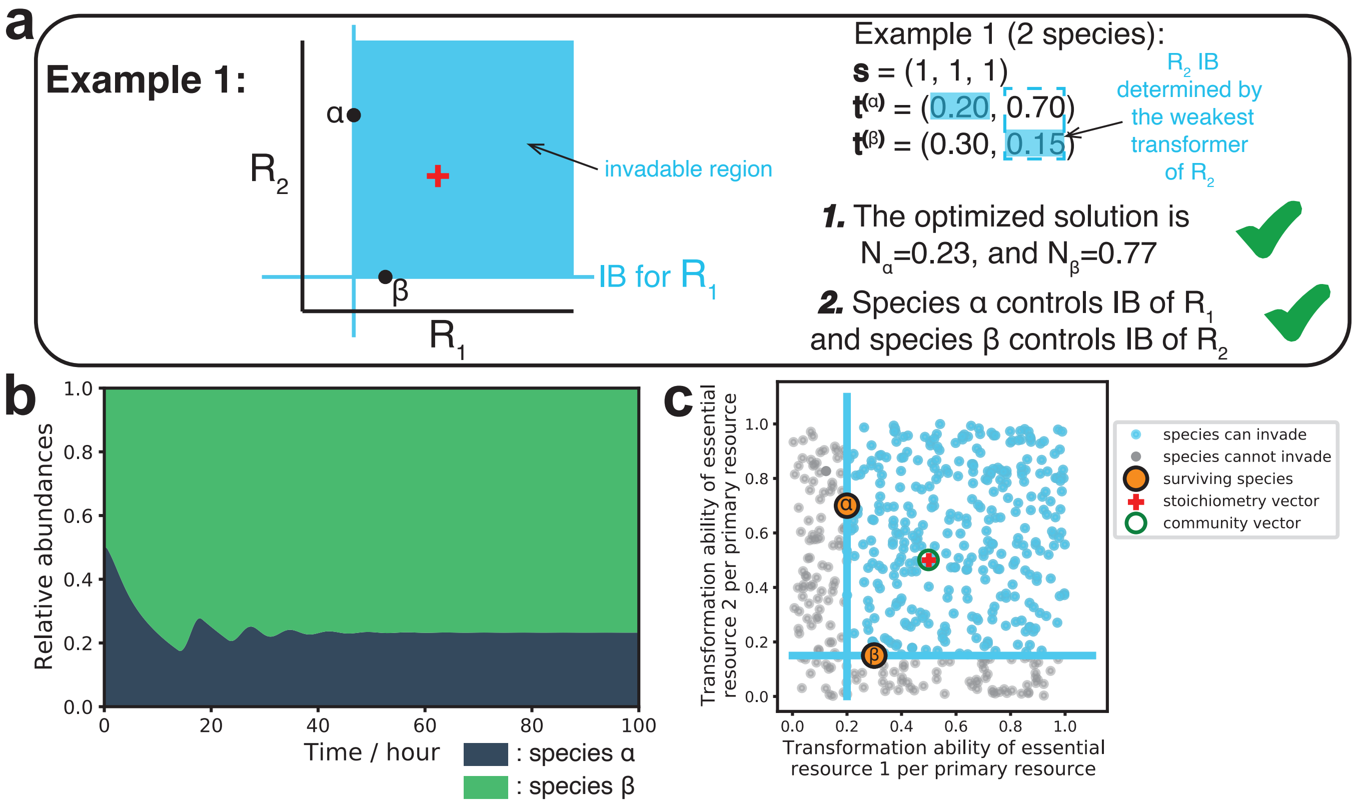


Supplementary Figure 3. **Agreement of abundance distribution and the invadable region between the graphical/algebraic predictions and model simulations when species' conversion efficiencies are different (i.e., $\sum_i t_i^{(\alpha)} \neq 1$).** **a** The graphical and algebraic approaches help to predict the stable steady state for a community with 2 species. **b** The simulation is performed in a chemostat environment, where the final abundance distribution for the stable steady state can be obtained by running the simulation with the initial 2 species. **c** Another randomly chosen species with a random transformation vector is generated and introduced into the stable steady state to see if it can invade the community. These invasion experiments are performed many times and the successful/failed invasions are colored light blue/grey.

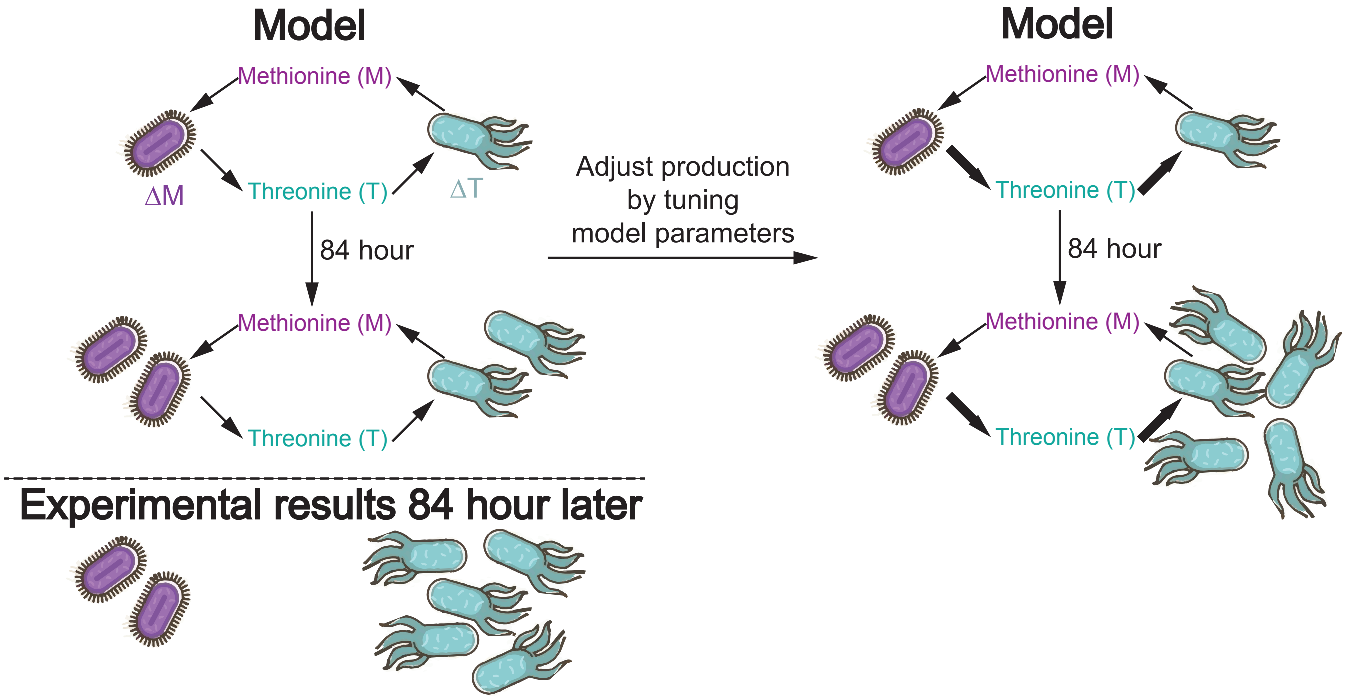


Supplementary Figure 4. **The iterative gradient-descent algorithm to fit the transformation abilities using fold-growth from pairwise co-culture experiments.** We consider the example of pairwise co-culture between $\Delta T$ and $\Delta M$. If the fold-growth of the strain $\Delta T$ predicted by our model exceeds or falls short of the experimental value, we adjust $\Delta M$'s transformation ability of $T$ upward or downward respectively to minimize the discrepancy.

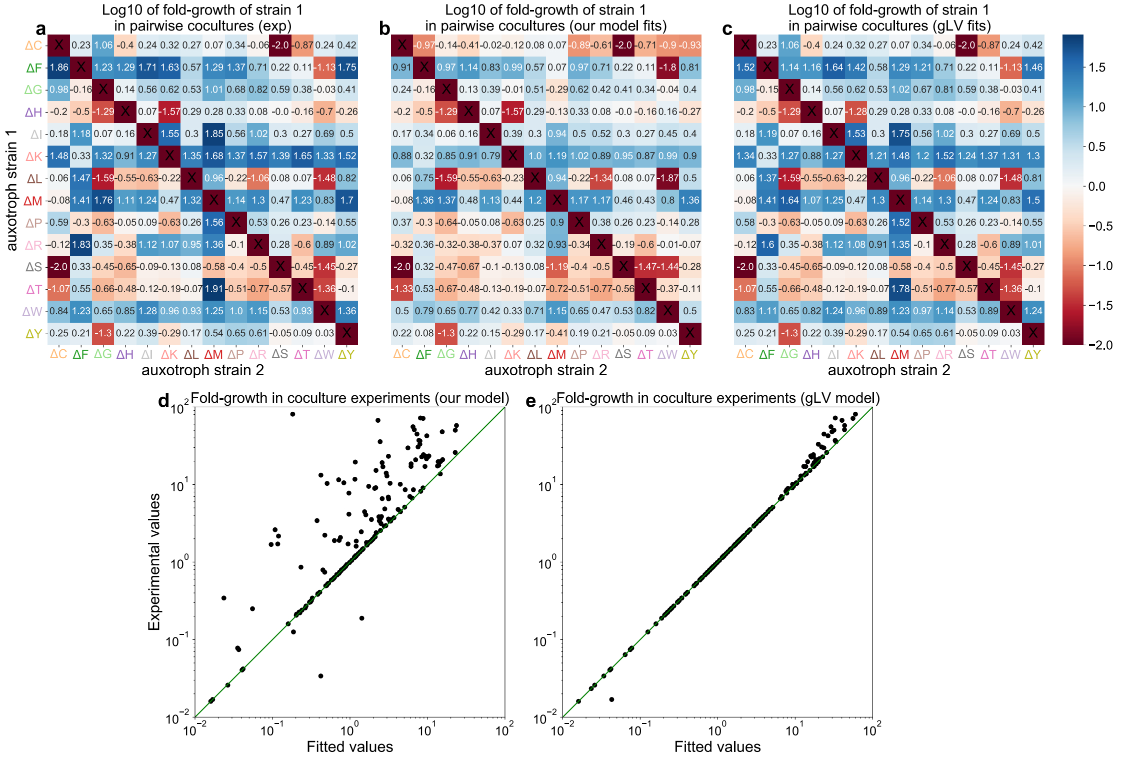


Supplementary Figure 5. **The gLV model overfits the experimentally measured data.** **a** The experimentally measured log10 of fold-growth of strain 1 in pairwise cocultures. b, Our model fitted log10 of fold-growth of strain 1 in pairwise cocultures. **b** Our model fitted log10 of fold-growth of strain 1 in pairwise cocultures. b, Our model fitted log10 of fold-growth of strain 1 in pairwise cocultures. **c** gLV fitted log10 of fold-growth of strain 1 in pairwise cocultures. b, Our model fitted log10 of fold-growth of strain 1 in pairwise cocultures. **d** The fitted values by our model and experimental values of fold-growth of each strain in co-culture experiments. **e** The fitted values by the gLV model and experimental values of fold-growth of each strain in co-culture experiments.

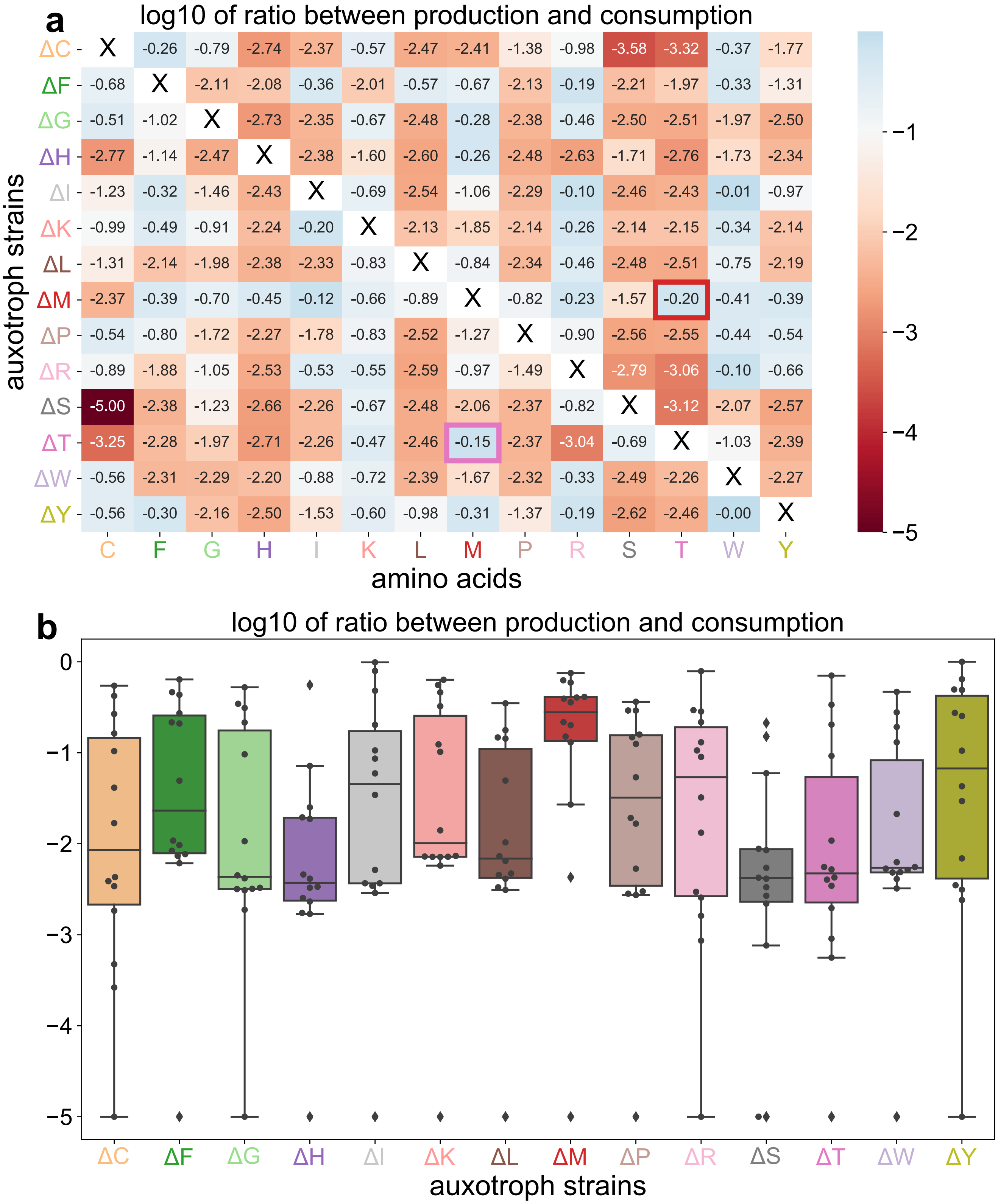


Supplementary Figure 6. **The log10 (logarithm with base 10) of the ratio between the best-fitted transformation vector and the assumed stoichiometry vector.** Before taking the log10, $10^{-5}$ is added to all values to avoid infinitely small numbers and to have a better visualization. **a** The heatmap of the log10 of the ratio between the best-fitted transformation vector and the assumed stoichiometry vector. **b** The boxplot of the log10 of the ratio between the best-fitted transformation vector and the assumed stoichiometry vector. All exact values are plotted as dots.

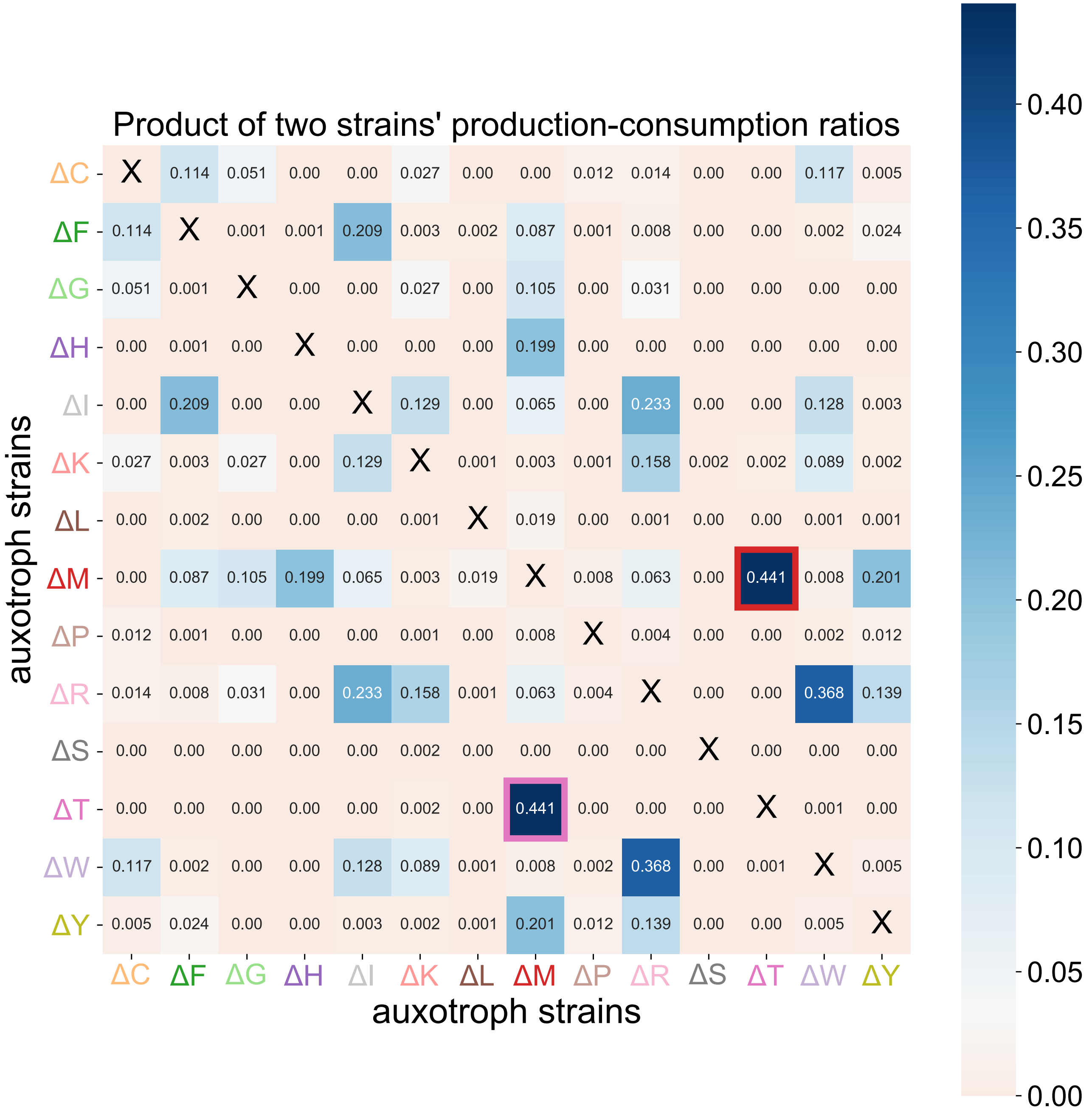


Supplementary Figure 7. $\Delta T$ **and** $\Delta M$ **form the best cooperative pair.** Color assignments for all strains and amino acids are consistent across all panels. The full names of amino acids are listed in the Methods section. The prefix $\Delta$ indicates a single amino acid auxotrophy (e.g., $\Delta T$ refers to the strain unable to synthesize Threonine ($T$)). For the element $\Delta X$-$\Delta Y$, its product of the two strains' production-consumption ratio is calculated by multiplying the transformation value of the strain $\Delta X$ to produce $Y$ with the value of the strain $\Delta Y$ to produce $X$.

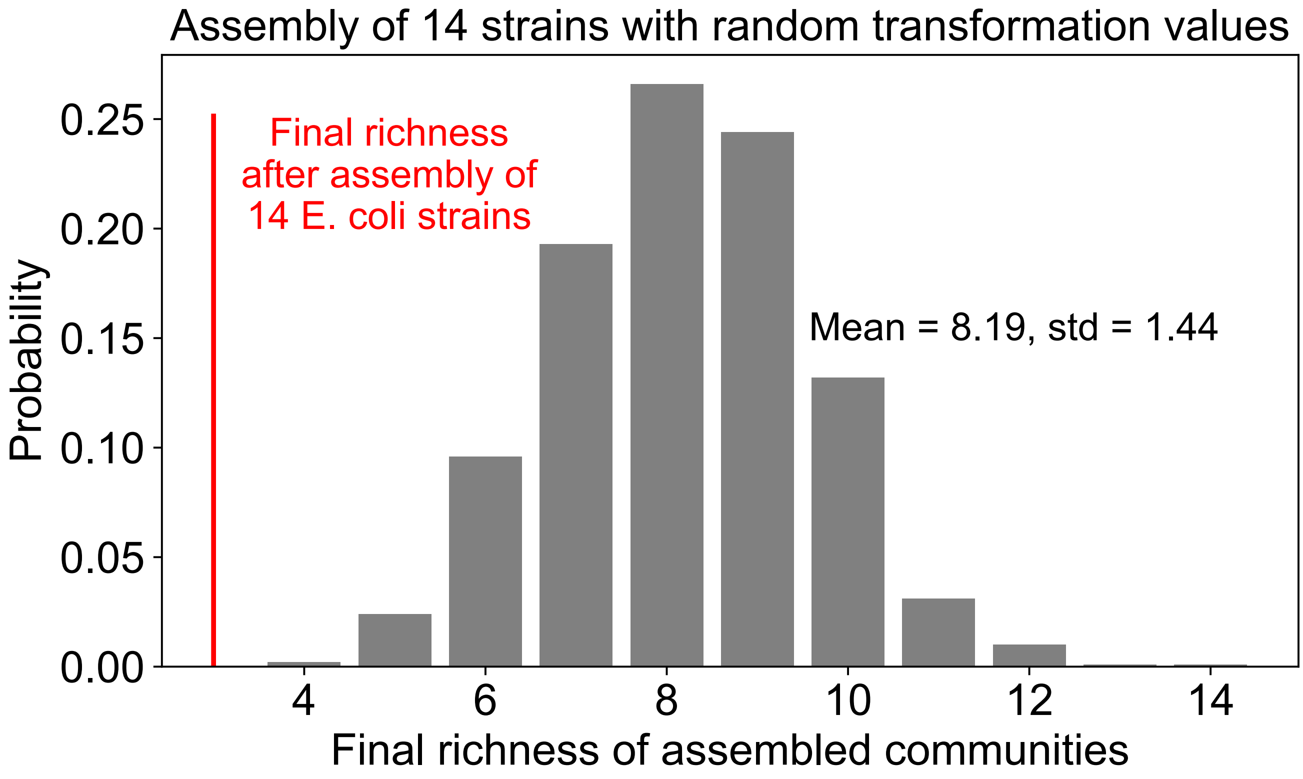


Supplementary Figure 8. **The assembly result of *E. coli* strains is less diverse than that of random strains.** 14 strains with randomly chosen transformation values are initialized with equal amounts to start the community assembly. Then the final richness of assembled communities is computed after simulating community assembly using our model. This type of random assembly is repeated 10,000 times to generate the probability distribution for the final richness of assembled communities.

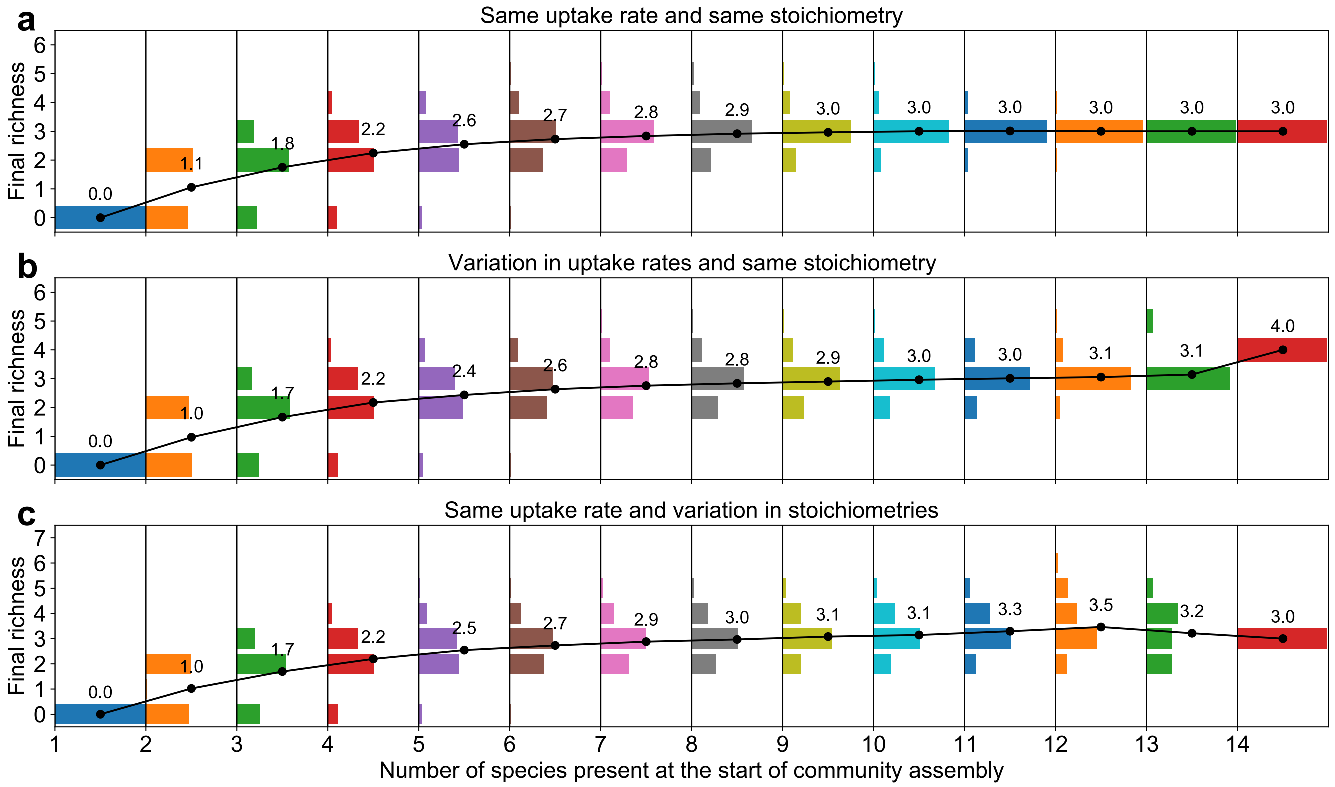


Supplementary Figure 9. **The variation in uptake rates or stoichiometries does not have a big impact on assembly results.** The normalized distribution of the final richness of assembled communities when different subsets of 14 single-resource auxotrophic species were initialized with equal amounts to start the community assembly. The distribution is generated for each initial richness based on all combinations of species. Dots connected by the black line represent the average final richness for each initial richness. The decimal numbers represent the value of the corresponding black dots. **a** The distribution of final richness of the assembled community when all species have the same uptake rate and stoichiometry (i.e., the synthetic *E. coli* strains). **b** The distribution of final richness of the assembled community when all species have different uptake rates and the same stoichiometry. The different uptake rates of the primary resource are created by multiplying the uptake rates in panel a by a random number drawn from a uniform distribution between 0.5 and 1.5. **c** The distribution of final richness of the assembled community when all species have the same uptake rate and different stoichiometries. The different stoichiometries are created by multiplying the stoichiometry in panel a by a random number drawn from a uniform distribution between 0.5 and 1.5.
